# Massively Parallel Screens Reveal Factors Influencing the Activity of a Diversity-Generating Retroelement Reconstituted in *Escherichia coli*

**DOI:** 10.1101/2025.08.18.670894

**Authors:** Irem Unlu, Marina K. Smiley, Vladimir Potapov, Yoan Renoux-Martin, Zhi-Yi Sun, Hoong Chuin Lim

## Abstract

Certain phages and microbes use Diversity-Generating Retroelements (DGRs) for targeted adaptation via a process called mutagenic retrohoming. This process relies on an error-prone reverse transcriptase that specifically misincorporates bases opposite adenines in the RNA template. The resulting mutagenized complementary DNA then integrates into a homologous target gene to introduce mutations at adenine residues. Although the functions of core DGR components are well-defined, we still don’t know how other cellular factors influence their activity. To address this, we reconstituted the DGR from the Bordetella phage BPP-1 in *Escherichia coli*. Our systematic analysis uncovered several cellular factors that influence its activity. By sequencing the cDNA, we discovered the reverse transcriptase is particularly mutagenic in the 5’-AAC-3’ and 5’-ACC-3’ contexts, which explains their abundance in natural DGR templates. A transposon sequencing screen identified both positive and negative regulators, including the single-stranded DNA exonuclease ExoI, which unexpectedly inhibits DGR activity through a nuclease-independent mechanism. Using a massively parallel reporter assay, we found that DGR preferentially edits targets located near the replication origin and co-directionally oriented with replication. In the preferred orientation, the target sequence is located on the lagging-strand template, where the extended single-stranded DNA gap likely facilitates base pairing with the incoming complementary DNA. Collectively, these findings advance our understanding of how DGRs function within the cellular context and enabled a 1000-fold improvement in editing efficiency, establishing a foundation for future targeted-mutagenesis applications.

## INTRODUCTION

Life constantly innovates through genetic trial and error. Organisms explore the vast genetic landscape by accumulating mutations to discover novel traits and adapt to changing environments. This exploration can be sped up through increased mutation rates, but these risks compromise genome integrity. Therefore, strategies enabling organisms to mutate rapidly but safely confer a profound evolutionary advantage, especially when a precise adaptation is urgently needed.

This delicate balance is exemplified by how our immune system generates antibodies to target foreign invaders. A vast array of unique antibodies is created by restricting heightened levels of DNA rearrangement and mutations to the antibody genes, sparing the rest of the cell’s DNA. Another distinct strategy for rapid, targeted adaptation involves diversity-generating retroelements (DGRs), found in phages, bacteria, and archaea (1–4). For example, the *Bordetella* phage BPP-1 employs DGR to selectively mutagenize adenines within the *mtd* (major tropism determinant) gene, which encodes the tail fiber protein responsible for host recognition (1). This generates a large repertoire of Mtd variants, promoting adaptation to the changing cell surface of its bacterial host (1).

DGRs diversify target genes through a process called mutagenic retrohoming (2). Extensive studies on the BPP-1 DGR have defined the operating principles of this mechanism (Fig. 1). Researchers have identified and characterized the dedicated reverse transcriptase (bRT) and its error-prone activity as the source of sequence diversity (5–9). They have also revealed the sequence elements crucial for target recognition and discrimination (5,6,10) and clarified how template RNA controls the initiation and termination of complementary DNA (cDNA) synthesis (8,9).

**Figure 1.**
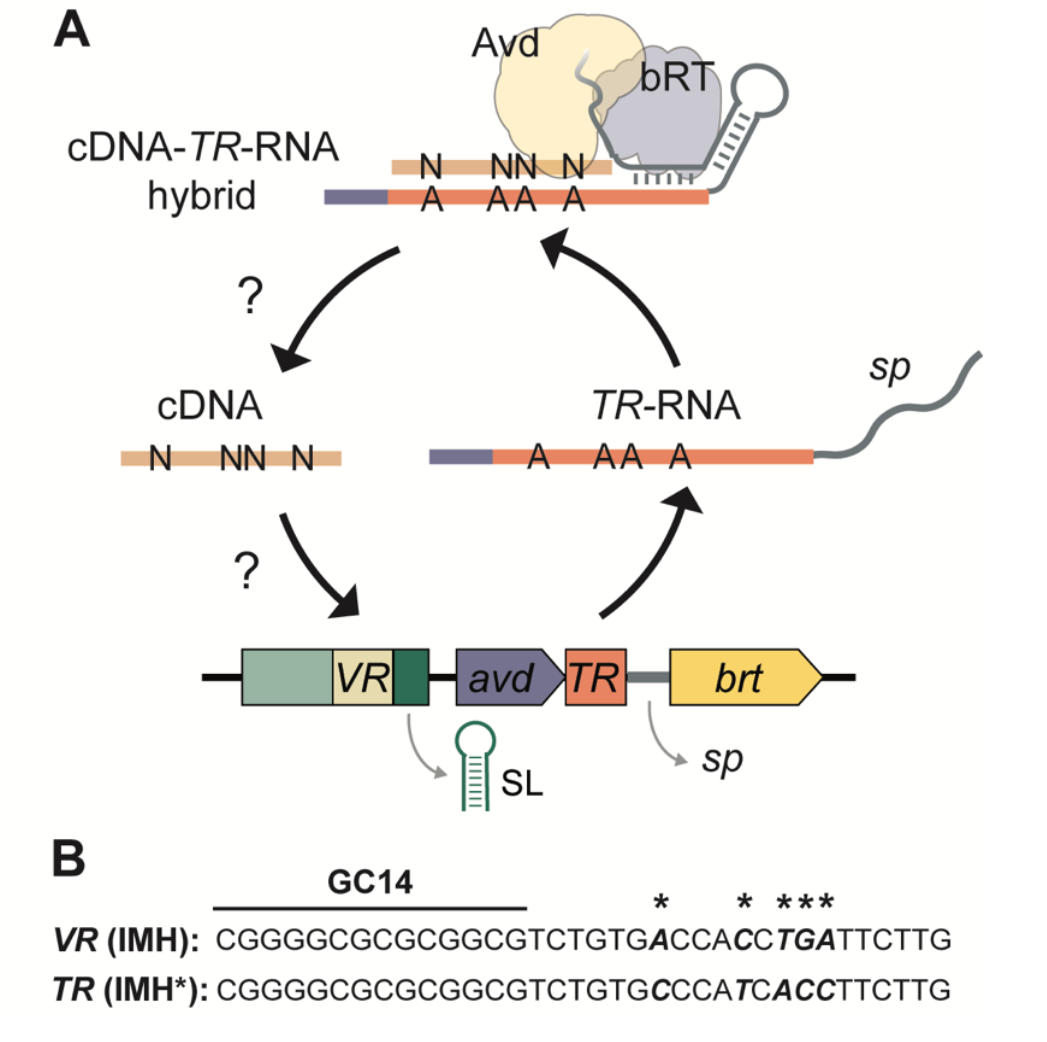
Mechanistic model of mutagenic retrohoming. **(A)** The BPP-1 DGR mediates mutagenic retrohoming using four core components: a reverse transcriptase (bRT), an accessory variable determinant (Avd), and two highly similar sequence repeats, namely the template repeat (*TR*) and variable repeat (*VR*) (2). The *TR*-RNA, which serves as the template, is flanked at the 5’ end by the coding sequence of *avd* and its 3’ by a spacer sequence (*sp*) located between *TR* and *brt*. This spacer sequence folds into an intricate structure that recruits the bRT-Avd complex (8). An internal adenine within this spacer then primes the initiation of cDNA synthesis within the 3’ junction of *TR* known as IMH* (6), resulting in a covalently attached cDNA-*TR*-RNA hybrid. A sequence motif within the 3’ end of the adjoining *avd* mRNA terminates reverse transcription (9). Due to the unique error-prone nature of bRT, the resulting cDNA contains mutations opposite template adenines (6,9). The precise mechanism by which this mutagenized cDNA then ‘retrohomes’ into the *VR* to modify the target gene remains incompletely understood. However, this process is known to be facilitated by two additional elements: 1) IMH located at the 3’ end of *VR*, which is nearly identical in sequence to IMH* (corresponding sequence in the BPP-1 DGR are shown in **B**) (1); 2) An essential inverted repeat that can form a stem-loop structure (SL) is located immediately 3’ of *VR* (10).

Despite this progress, several aspects of this process remain incompletely understood. For example, we lack a mechanistic understanding of how the resulting cDNA is incorporated into the target gene. It is also unknown how DGR functions within a cellular environment, including whether its activity is influenced by host factors or coordinated with other cellular processes. A key unanswered question is whether DGRs can function in tractable model organisms like *Escherichia coli* and *Bacillus subtilis*, which do not naturally carry DGRs (11–13).

Here, we report the successful reconstitution of the mutagenic retrohoming activity of the BPP-1 DGR in *E. coli*. Using a combination of deep sequencing, massively parallel reporter assays, and genetic analysis, we studied this reconstituted DGR system. Our findings revealed how nucleotides surrounding adenines in the RNA template affect the error profile of cDNA and identified cellular factors that influence DGR activity. We also implicated DNA replication as a key driver of cDNA retrohoming. Our work significantly enhances the understanding of how DGRs function inside a cell. By leveraging these new insights, we were able to optimize the system, improving DGR-mediated gene editing by more than three orders of magnitude. This advance paves the way for more efficient characterization and biotechnological application of DGRs in a tractable organism.

## RESULTS

### Reconstitution of the BPP-1 DGR in *E. coli*

We set out to reconstitute the archetypal BPP-1 DGR in *E. coli*, leveraging the extensive genetic tools available in this bacterium to better understand the mechanism of DGR. We encoded the essential BPP-1 DGR components on two plasmids (Fig. 2A): pDGR0 carried the *avd-TR-brt* operon under an arabinose inducible promoter, while pTarget contained a target gene immediately followed by the essential regulatory elements: GC14, IMH, and stem-loop.

**Figure 2.**
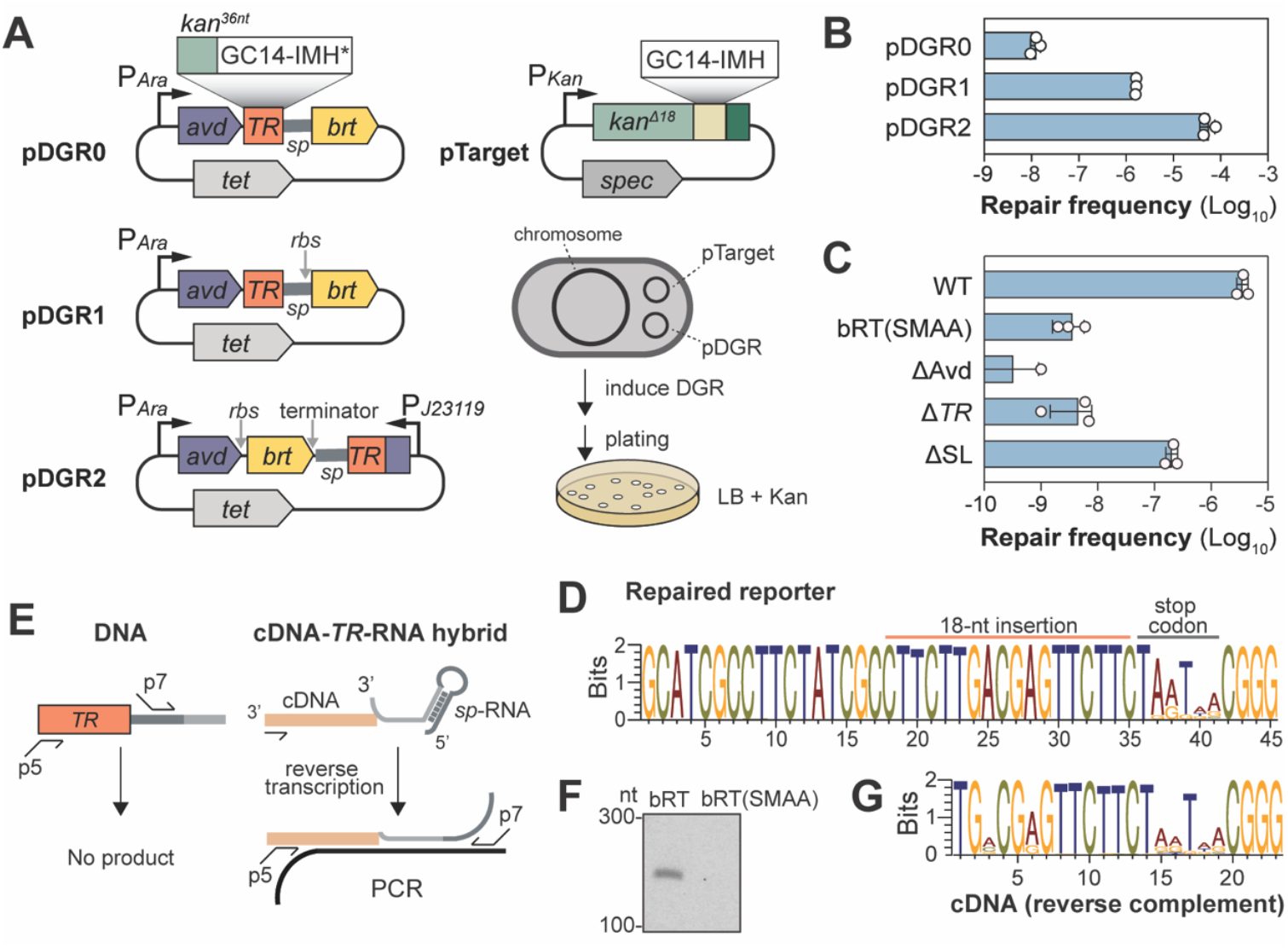
Reconstitution of BPP-1 DGR-mediated editing in *E. coli*. **(A)** Schematic of the DGR-mediated kanamycin repair reporter assay (15). Left, Different pDGR variants, which encode all essential DGR components except for the target. The *TR* consists of the last thirty-six nucleotides of the *kanR* gene, followed by tandem stop codons (TGATAA). Right, pTarget encodes the target gene, which is a defective *kanR* gene terminated by TAATAA. DGR-mediated repair of the reporter was monitored by quantifying the frequency of kanamycin-resistant clones (repair frequency). **(B)** Bar graph showing the repair frequency for pDGR0, pDGR1, and pDGR2 in *E. coli*. Each of these plasmids was used in tandem with pTarget to transform *E. coli* strain MG1655. Freshly transformed colonies were used to inoculate LB+Tet10+CM15. The cultures were grown for 1.5 hours at 37°C, after which arabinose was added to induce expression. The cultures were then grown for an additional 4.5 hours. Finally, the cells were then plated on rich media agar supplemented with KAN25 to select for resistant clones. See Methods for details. **(C)** Bar graph comparing the repair frequency of pDGR1 and variants with the indicated mutations and deletions. These include: bRT(SMAA), YMDD>SMAA active site mutations in *brt*; Δa*vd*, three nonsense mutations in *avd*; Δ*TR, TR* deletion; ΔSL, stem-loop deletion. Data are mean ± standard deviation (n = 3 biological replicates). **(D)** Weblogo showing the recovery of the missing 18-nt in pTarget upon induction of DGR expression. Plasmids were harvested from a pool of kanamycin resistant clones containing both pTarget and pDGR1. The plasmids were then used as a template, and the target region was amplified by PCR for deep sequencing. Only sequencing reads showing an 18-nucleotide insertion in the reporter were used to generate the Weblogo. **(E)** Scheme for specifically amplifying the cDNA-*TR*-RNA for deep sequencing while avoiding DNA template amplification. **(F)** Agarose gel showing the RT-PCR product is dependent on an intact bRT. **(G)** Weblogo showing the results of cDNA sequencing. The reverse complement of the cDNA sequence is shown to facilitate easy comparison to the reporter in panel **D**. This sequence is shorter because the primer binding region is excluded.

To assess DGR activity, we used a reporter system previously used to characterize the BPP-1 DGR in its native host (10). In this assay, the target gene was a defective kanamycin resistance cassette lacking the sequence for its last six amino acids. The *TR* was programmed with the sequence encoding the final twelve amino acids of the full-length cassette (Fig. 2A). This design should enable DGR to transfer the missing sequence to the target through a cDNA intermediate, thereby “repairing” the defective kanamycin resistance gene. We then measured DGR activity by quantifying the frequency of kanamycin-resistant clones, which we termed the repair frequency.

Our initial experiments to reconstitute DGR activity in *E. coli* MG1655 using pTarget and pDGR0 yielded kanamycin-resistant clones at a meager ~10^−8^ frequency (Fig. 2A-B). Absence of DGR function was likely due to the lack of a clear ribosomal binding site (RBS) upstream of the *brt* gene within the native BPP DGR operon. Introducing a consensus *E. coli* RBS before *brt* in the new construct (pDGR1) resulted in kanamycin-resistant colonies (Fig. 2B). As in the native host, the gain of kanamycin resistance was not driven by homologous recombination between the *VR* and the *TR*. Deleting the major recombinase gene, *recA*, only minimally affected the repair frequency (Fig. S1). Instead, resistance arose only with intact essential DGR components (Fig. 2C). Inactivating any essential DGR component—bRT, Avd, or *TR*—reduced the frequency of kanamycin resistant clones to ~10^−9^ (Fig 2C). The only exception was the inverted repeat found downstream of IMH. This inverted repeat has stem-loop forming capability and is essential for BPP-1 DGR activity in its native *Bordetella* host (10). In *E. coli*, however, this inverted repeat is dispensable; we observed significant repair frequency (ten-fold above background) when it is deleted from pDGR1, despite a 22-fold reduction (Fig. 2C). We concluded that this reconstituted DGR activity recapitulates important features found in the native host.

While we successfully reconstituted DGR activity in *E. coli*, the observed repair frequency of 10^−6^ was quite low. To enhance this, we created pDGR2 (Fig. 2A). In this construct, we removed the *TR-*RNA from the operon and placed it under the control of P_J23119_, a strong constitutive synthetic promoter often used for expressing single-guide RNAs in CRISPR applications (14). This modification improved repair frequency by 35-fold compared to pDGR1 (Fig. 2B). However, the increased repair frequency appeared to result from higher bRT levels—possibly a side effect of the genetic rearrangement—rather than higher *TR-*RNA levels (Fig. S2A-B).

DGR activity was also evident at the DNA level. Our analysis showed that 59% (pDGR1) and 43% (pDGR2) of the *VR* regions have recovered the missing 18 nucleotides (Fig. S3A). We believe these percentages reflect the proportion of the 20-40 copies of pTarget in each cell that were restored by DGR, rather than indicating that roughly half of the clones contained repaired reporters. Supporting this, Sanger sequencing of plasmids isolated from individual clones often showed heterogeneity in this region (Fig. S3B). By contrast, among roughly 50 spontaneously kanamycin-resistant clones from our mutant bRT control (arising at a frequency of <10^−8^), only 3-4% had recovered the missing sequence (Fig. S3A). This suggests that suppressor mutations located elsewhere in the genome, instead of recombination between *TR* and *VR*, mainly contribute to background resistance.

This analysis also identified the characteristic A-to-N mutations within the restored reporter, but only in the four adenines in the stop codons, not in the coding region (Fig. 2D). We confirmed this was not due to target incorporation efficiency for cDNAs with the coding region mutations, as programmed synonymous mutations at non-adenine positions were incorporated at 100% efficiency without affecting the repair frequency (Fig. S4A-B). Furthermore, direct sequencing of the cDNA showed significant misincorporation opposite coding adenines, indicating that mutagenesis was not suppressed during reverse transcription (Fig. 2E-G). We conclude that missense mutations in coding adenines inactivated the reporter gene and were thus selected against. Consequently, the observed repair frequency underestimated the true frequency of DGR-mediated target editing.

Together, these results demonstrated that DGR’s characteristic mutagenic retrohoming activity can be faithfully reconstituted in *E. coli* with its core components alone.

### Sequence context biases nucleotide incorporation for A-to-N mutations

Our cDNA sequencing analysis (Fig. 2G) confirmed a previous in vitro finding that misincorporation choice and rate vary depending on the adenine’s position within the RNA template (7). This suggests that the surrounding sequence context influences the error-prone reverse transcription; however, this relationship has not been systematically characterized. To address this, we designed a pDGR2 variant with a new template encoding all 16 possible (NAN) trinucleotides within a single *TR* RNA. To control for positional effects, we created two additional variants where the template was split into three parts and rearranged to assess how being at the front, middle or the end of the RNA template affected mutagenesis (Fig. 3A; *TR1, TR2*, and *TR3*). We then expressed these RNAs along with bRT and Avd in *E. coli*, and used deep sequencing to profile the resulting cDNAs.

**Figure 3.**
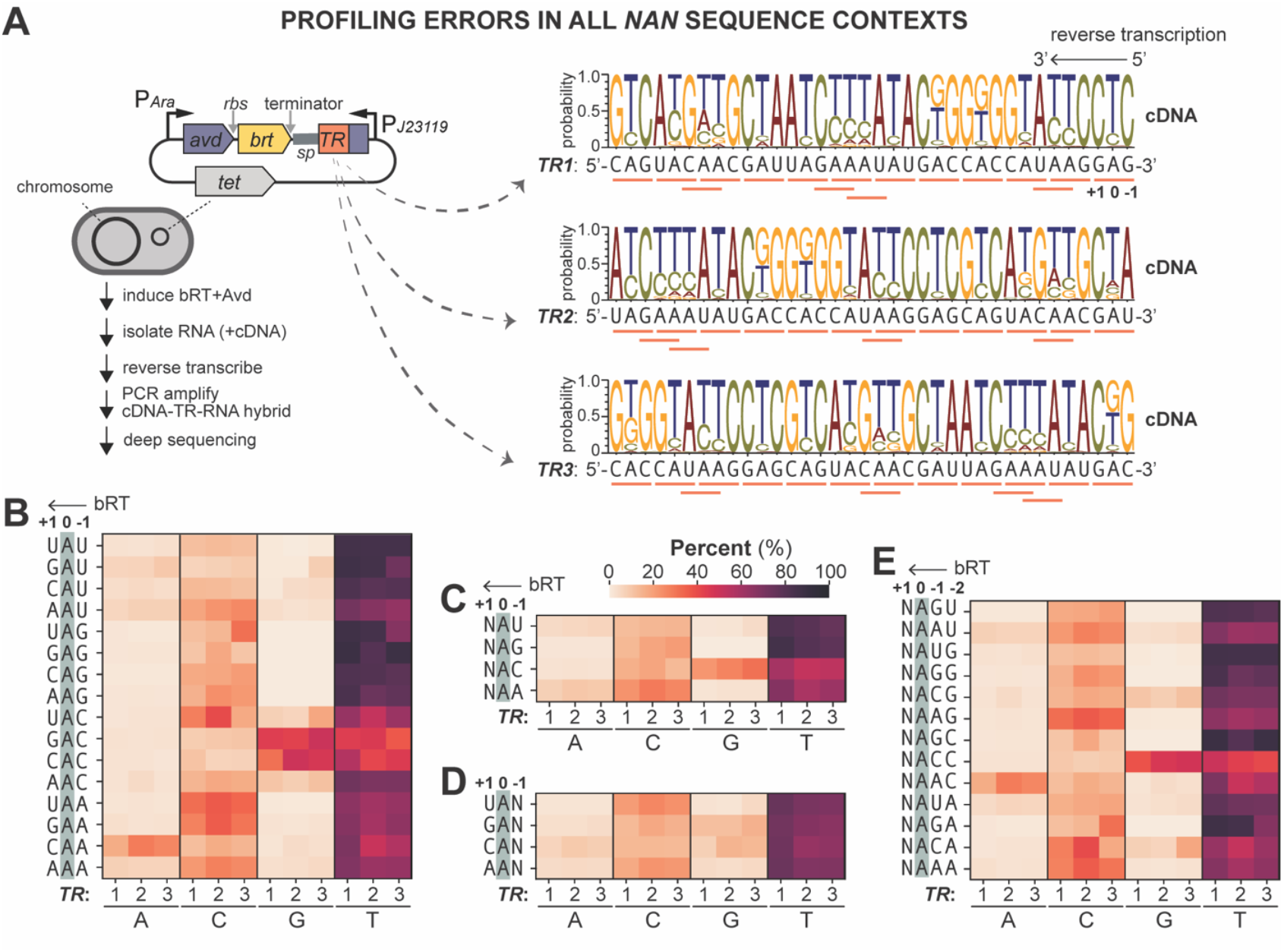
Effect of sequence context on the fidelity of BPP-1 bRT. **(A)** Left, Scheme for cellular profiling the mutational signature of bRT across various template sequence contexts. Right, pDGR2 was programmed with *TR1, TR2*, and *TR3*, each containing all 16 possible NAN triplets (underlined in orange) in different order. Weblogos display the error profiles of cDNA synthesized by bRT from *TR1, TR2* and *TR3* RNA templates. **(B)** Heatmap shows the percentage of A, C, G and T incorporation opposite template adenines within all 16 NAN contexts, using *TR1, TR2* and *TR3* as RNA templates. The Y-axis shows the RNA sequence contexts. **(C)** Heatmap shows the percentage of base incorporations opposite template adenines in the N<u>A</u>U, N<u>A</u>G, N<u>A</u>C and N<u>A</u>A contexts, highlighting the effect of the nucleotide immediately 3’ (−1 position) of the adenine. **(D)** Heatmap shows the percentage of base incorporations at template adenines in the U<u>A</u>N, G<u>A</u>N, C<u>A</u>N and A<u>A</u>N contexts, highlighting the influence of the nucleotide immediately 5’ (+1 position) of the adenine. **(E**) Heatmap shows the percentage of base incorporations opposite template adenines in the indicated contexts, highlighting the combined influence of nucleotides at both the −1 and −2 positions relative to adenine. Data in all heatmaps represent the average of biological triplicates for each *TR*.

Our analysis revealed that the mutational profile remained largely consistent across the three RNA templates (Fig. 3B), indicating that the specific position of adenines within the RNA template had minimal impact. However, we uncovered notable differences when comparing our findings to previous *in vitro* measurements (7). Specifically, the average misincorporation rate of 30.65% across all trinucleotide contexts in *E. coli* is lower than the reported 50% rate measured *in vitro* (Fig. 3B) (7). Furthermore, cytosine (C) misincorporation predominated (18.2%) in cells, which contrasts with the reported preference for adenine (A) misincorporation *in vitro* (7). These discrepancies likely stem from an overrepresentation of specific sequence motifs that bias adenine misincorporation in the RNA template used for the previous *in vitro* analysis (see below and Discussions).

Further analysis uncovered interesting context dependence in bRT-mediated cDNA mutagenesis. We observed that bases immediately 3’ of template adenines (at position −1) influenced the mutagenesis profile more than those immediately at the 5’ end (+1) (Fig. 3C-D). A prominent example of this context dependence was seen with adenines in the 5’-N<u>A</u>C-3’ context (where N is any base at position +1, and C is at −1) (Fig. 3C). Specifically, guanosine (G) misincorporation was markedly elevated (26.8%) in this context, compared to N<u>A</u>A (2.4%), N<u>A</u>G (0.08%) and N<u>A</u>T (2.5%) contexts (Fig. 3C). Even more strikingly, G misincorporation was higher in the G<u>A</u>C (45.9%) and C<u>A</u>C (42.6%) contexts (Fig. 3B). This heightened G misincorporation could be caused by the presence of G and C at +1 position, or potentially the presence of an additional cytosine at the −2 position: C<u>A</u>C(C) and G<u>A</u>C(C) (Fig. 3E), as opposed to A<u>A</u>C(G) and T<u>A</u>C(A) (Fig. 3A). We also found bRT was more prone to misincorporate adenine instead of the more common error, cytosine, within the 5’-<u>A</u>AC-3’ context (Fig. 3E). This clarifies why significant adenine misincorporation occurred at C<u>A</u>A(C), but not in the T<u>A</u>A(G), G<u>A</u>A(A) and A<u>A</u>A(T) contexts (Fig. 3A-B). Altogether, our systematic analyses characterized the error-profile of bRT when transcribing template adenines across diverse sequence context, pinpointing several instances of altered enzyme behavior.

### Multiple cellular factors influence DGR activity

Having reconstituted DGR and characterized its mutagenic properties in *E. coli*, we next tried to improve upon its editing efficiency (~10^−5^), which is too low for practical applications. We reasoned that a deeper understanding of how DGR functions within the cell would help inform strategies for improving efficiency. To achieve this, we conducted a high-throughput genetic screen using transposon-sequencing (Tn-seq) (16) to identify cellular factors influencing DGR activity.

To streamline the screen, we integrated the reporter cassette directly into the *E. coli* genome. Among the four tested chromosomal loci (Fig. 4A), we chose reporter insertion at the 317° locus (strain HCL26), which exhibited the highest DGR-mediated repair frequency, for our screen. We mutagenized HCL26 using a mariner-based transposon, generating a transposon mutant library of about 150,000 unique clones (Fig. 4B). This library was then transformed with pDGR2 to initiate reporter repair. To quantify the repair frequency of all mutants simultaneously, we deep sequenced the transposon distribution before (Input) and after kanamycin selection (Output). In principle, the read counts of each transposon insertion should be proportional to its abundance within the library. Therefore, genes with enriched transposon insertions in the Output relative to the Input are considered potential inhibitors of DGR, as their disruption would increase reporter repair. Conversely, transposon insertions in genes that promote DGR activity would impair reporter repair and become depleted in the Output.

**Figure 4.**
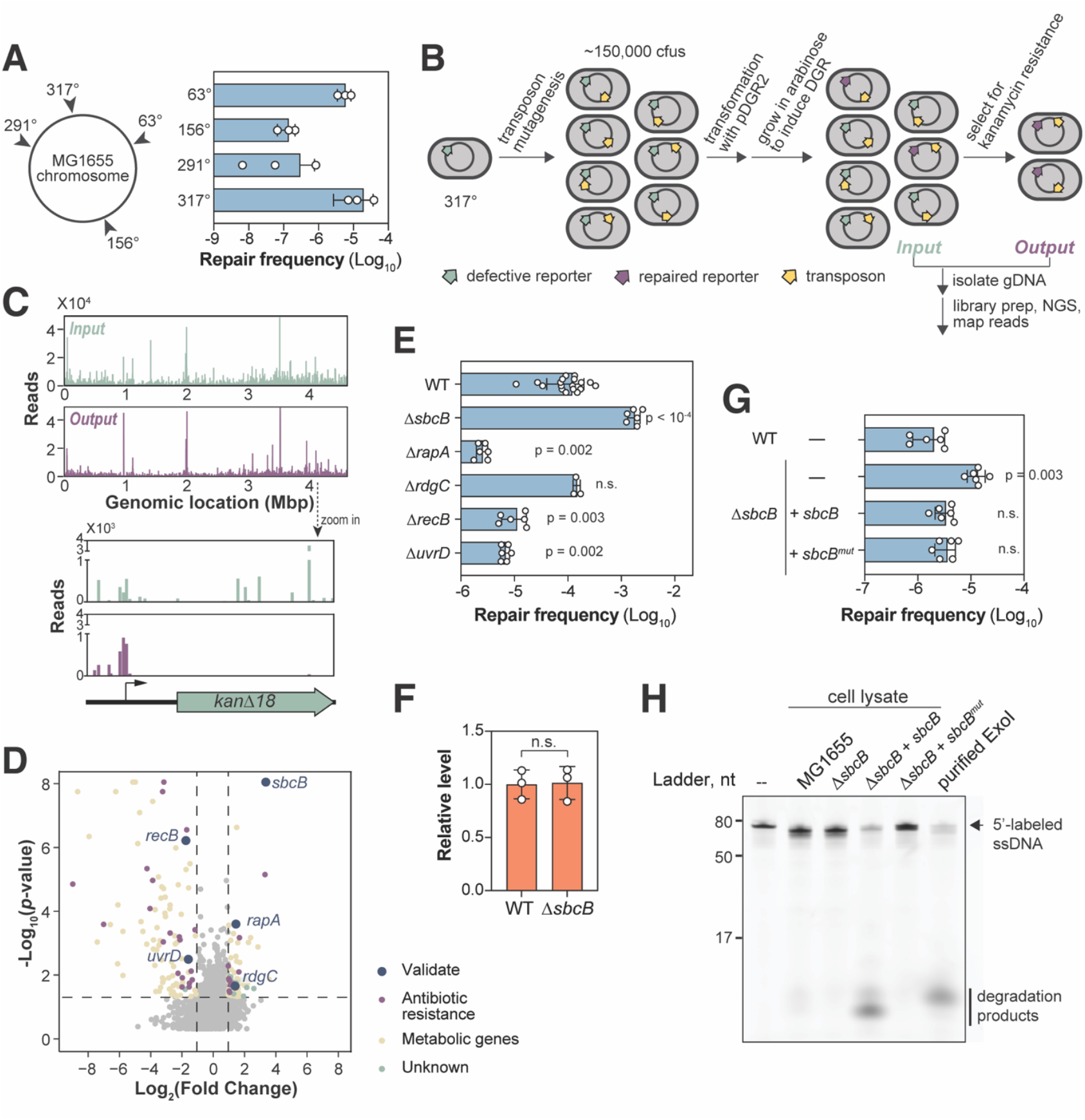
Screening for cellular factors affecting BPP-1 DGR activity. **(A)** Left, Chromosomal locations of reporter insertion. Right, Bar graph comparing the repair efficiency of reporters at the indicated chromosomal locations in MG1655. *E. coli* strains used in this experiment were: 63°(HCL34), 156°(HCL19), 291°(HCL24), and 317°(HCL26). **(B)** Schematic of Tn-seq workflow to identify factors affecting DGR-mediated reporter repair. See Methods for details. **(C)** Normalized Illumina read counts from each transposon insertion of the Input and Output libraries. Bump out shows changes in transposon distribution within the reporter cassette in the Input and Output libraries. **(D)** Volcano plot of transposon sequencing results. Vertical dotted lines represent thresholds for 2-fold increase or decrease in read counts. Data above the horizontal dotted line have Mann-Whitney *p*-value of 0.05 or less. **(E)** Bar graph showing repair frequency of a chromosomal reporter in *E. coli* with the indicated genetic background. All mutants—Δ*sbcB* (HCL84), Δ*rapA* (HCL102), Δ*rdgC* (HCL144), Δ*recB* (HCL138), and Δ*uvrD* (HCL137)—were derivatives of HCL26 (WT). Error bars represent standard deviation of at least biological triplicates. All p-values (unpaired Student’s t-test) were calculated by comparing each deletion mutant to the MG1655 control. n ζ 3 biological replicates. **(F)** Bar graph showing the relative abundance of *TR*-RNA-cDNA hybrid in MG1655 vs Δ*sbcB* (HCL84) strain as measured by qRT-PCR. n = 3 biological replicates **(G)** Bar graph comparing the repair frequency of a chromosomally integrated reporter in MG1655 (WT) or Δ*sbcB* (HCL84) strain. These strains also contained either an empty vector pNEB309 (–), pNEB310 (expressing *sbcB*) or pNEB311 (expressing the nuclease-deficient *sbcB*^*Mut*^). **(H)** Cell-lysate-based ssDNA nuclease assay. A 5’ Alexa488 fluorescently labelled ssDNA (see Table S3 for sequence) was incubated for 10 minutes at room temperature with lysates prepared from strains used in **(G)** or commercially available purified ExoI (NEB MS293). These samples and the unreacted ssDNA substrate (–) were resolved on a 15% polyacrylamide gel before fluorescence imaging.

This analysis revealed a set of genes, notably *envZ* (encoding a stress response kinase), whose transposon insertions showed disproportionately high sequencing reads in both the Input and Output (Fig. 4C). Their overrepresentation likely stemmed from increased resistance towards chloramphenicol (Fig. S5). Furthermore, our analysis identified 175 genes with at least twofold difference in representation between the Input and Output (p-value < 0.05). Among these, 97 genes, when interrupted by transposon insertion, became depleted in the Output (Fig. 4D). These included the reporter cassette (Fig. 4C), and genes previously found to help *E. coli* survive kanamycin exposure (Table S1), confirming the effectiveness of our screen. This list also included many genes encoding metabolic function (Table S1), the significance of which was not investigated further in this study.

We selected five hits involved in RNA or DNA metabolism—*sbcB* (*xonA*), *rapA, rdgC, recB*, and *uvrD*—for validation. Deleting each of these from the HCL26 strain confirmed several screen findings (Fig. 4E): *recB* and *uvrD* deletions reduced repair frequency by 11- and 18-folds, respectively. Conversely, deleting *sbcB* increased repair frequency by 15-fold. While *rdgC* deletion had no effect, *rapA* deletion surprisingly decreased repair frequency, contradicting our Tn-seq finding. These results indicate that RecB, UvrD and RapA promote DGR activity, while ExoI (encoded by *sbcB*) inhibits it.

### ExoI inhibits DGR independent of its exonuclease function

We focused our characterization on ExoI, a nuclease that degrades single-stranded DNA (ssDNA) from the 3’ to 5’ end, to understand how it inhibits DGR editing. We first confirmed that the increased repair frequency in the Δ*sbcB* strain was not due to an elevated kanamycin resistance or activation of the RecF recombination pathway (17) (Fig. S6). We then considered if deleting ExoI might simply stabilize cDNA. To directly test this, we compared the cDNA levels between the wild-type and Δ*sbcB* strains using quantitative PCR (qPCR). Our amplification strategy specifically targeted the cDNA-*TR*-RNA hybrid (Fig. 2E-F), which is thought to be the cDNA species that integrate into the target (6). This analysis revealed that the hybrid level remained unchanged in the Δ*sbcB* strain (Fig. 4F), suggesting that hybrid stabilization does not account for the heightened DGR activity in cells lacking ExoI. Consistent with this, our Tn-seq data indicated that inactivating any nuclease other than ExoI did not enhance repair frequency (Fig. S7). This is particularly noteworthy because deletion of ExoI, along with other *E. coli* nucleases like ExoVII, ExoX and RecJ, has been shown to boost recombineering, presumably by stabilizing the ssDNA substrate (18).

Given this, we explored the possibility that ExoI might inhibit DGR through a nuclease-independent function, which has precedence both in *E. coli* (17,19–21) and *Vibrio cholerae* (22). Using a plasmid, we expressed either wild-type ExoI or ExoI^Mut^, a nuclease-deficient variant with D15A and E17A active site mutations, in the Δ*sbcB* background. Expression of both ExoI and ExoI^Mut^ in trans suppressed the increased repair frequency in the Δ*sbcB* strain equally (Fig. 4G). Thus, the nuclease function of ExoI is dispensable for inhibiting DGR.

To confirm that ExoI^Mut^ was indeed nuclease-defective, we used a cell-lysate-based nuclease assay (22). Cell lysates were prepared from the strains used in Fig. 4G. We then monitored the degradation of a 5’-labeled ssDNA substrate in the lysates. In this assay, the ssDNA substrate remained largely intact in lysates from MG1655 and *ΔsbcB* cells (Fig. 4H). In contrast, lysates from *ΔsbcB* cells expressing wild-type ExoI from a plasmid exhibited robust ssDNA degradation, similar to commercially available ExoI. Expression of ExoI^Mut^ did not show this activity, confirming its nuclease deficiency. These results reinforce the qPCR finding (Fig. 4F), indicating that ExoI normally suppresses DGR activity through another function unrelated to its exonuclease function.

Beyond confirming the deficiency of ExoI^Mut^, the nuclease assay also revealed significantly higher exonuclease activity in the lysate when ExoI was expressed in trans (Fig. 4H, compare Lane 2 and 4). However, this heightened exonuclease activity did not reduce repair frequency below that of the wild-type strain (Fig. 4G). This is notable because it indicated that the inhibitory effect of ExoI is saturable, which further argues against an inhibitory mechanism involving a catalytic function. Instead, inhibition is likely mediated through a stoichiometric interaction between ExoI and a yet unknown binding partner. Because ExoI^Mut^ inhibits DGR, it must still retain this target binding capacity.

### Chromosomal location and orientation of target genes affect DGR editing efficiency

In our characterization of chromosomal targets (Fig. 4A), we observed a significant variation in the repair frequency among four genomic targets, differing by as much as 190-fold. To investigate whether target orientation also influences DGR editing, we constructed four more reporter strains, in which the orientation of the target at those chromosomal loci was reversed. Reversal of target orientation had a notable effect on repair frequency across all four loci, with the two reporters at the 291^°^ locus showing the highest difference of 44-fold based on their orientation (Fig. 5A).

**Figure 5.**
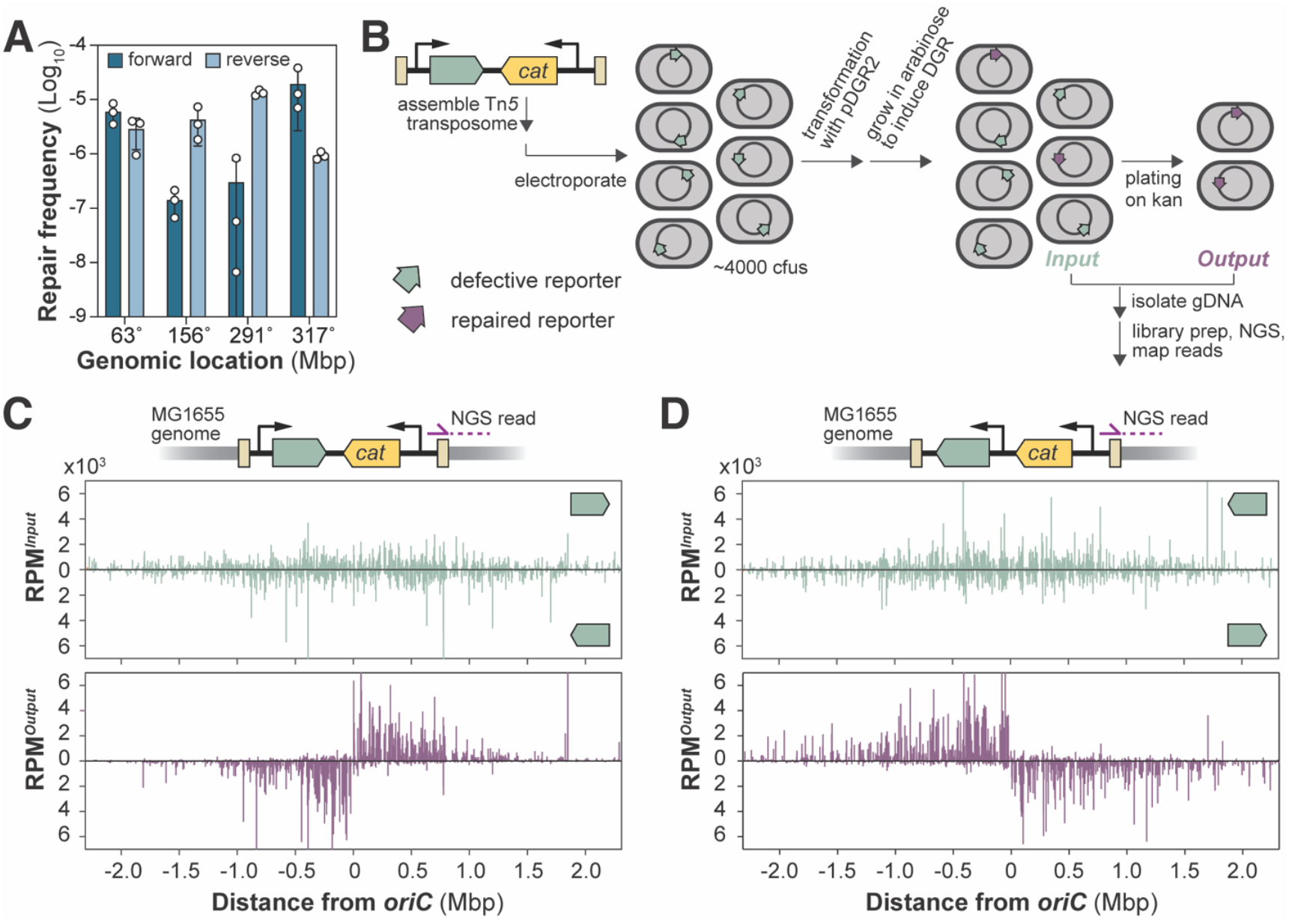
Both chromosomal location and target gene orientation influence DGR activity. **(A)** Bar graph comparing the repair efficiency of reporters inserted at the indicated chromosomal locations in both orientations in *E. coli*. DGR components were expressed from pDGR2. n = 3; error bars indicate standard deviation. **(B)** Schematic of a high-throughput workflow for profiling the efficiency of DGR-dependent editing of thousands of chromosomally integrated reporters at once. **(C)** Schematic showing the reporter orientation within the transposon. Transposon sequencing workflow was used to map the right transposon-chromosome junction. In this transposon design, the reporter is co-directional with the sequencing read. Normalized Illumina read counts (reads per million, RPM) of mapped transposon insertion sites across the MG1655 genome for both the Input and Output libraries. Note that the genomic coordinate is adjusted to center the data around *oriC* (3,925,797 bp). **(D)** Same as described in **(C)**, except that the reporter orientation is reversed in the transposon used in this experiment. Consequently, the reporter orientation is contra-directional with the sequencing read.

To identify the optimal genomic locus in an unbiased manner and determine whether a genome-wide trend exists, we simultaneously measured the repair frequency of reporters at roughly 5000 chromosomal loci (Fig. 5B). These reporters were randomly inserted into the MG1655 chromosome using the Tn*5* transposase. The resulting reporter strains were then pooled, and their repair frequencies were measured in parallel using the Tn-seq workflow.

Our analysis identified about 5000 unique target insertions throughout the chromosome in both orientations in the starting library (Fig. 5C, top). However, reporter distribution shifted dramatically after selecting for kanamycin resistance to enrich for DGR-restored reporters (Fig. 5C, bottom). This altered representation validated the genome-wide effect of both reporter location and orientation, as seen in our previous experiment (Fig. 5A). Reporters located near the origin of replication (*oriC*) became overrepresented, indicating higher repair frequency in this region. Beyond this positional effect, reporter orientation profoundly impacted their representation across the genome (Fig. 5C). On the left arm of the chromosome, reporters oriented leftward were enriched, whereas left-facing reporters on the right arm were specifically depleted. This striking pattern indicates that the favored orientation abruptly switches at *oriC*, consistently preferring reporters that point away from the origin of replication.

We confirmed that the observed trends were solely due to target orientation and not the transposon. By reversing the reporter orientation in a new transposon design, we observed a corresponding reversal in the orientations of preferentially repaired transposons (Fig. 5D). This indicated consistent bias in the preferred target orientation, regardless of the transposon design (Fig. 5C-D). Therefore, DGR editing efficiency is determined by a combination of its genomic position relative to *oriC* and target orientation.

### Replication directionality dictates the preferred target orientation

In *E. coli*, DNA replication initiates at the *oriC*, forming two replication forks that migrate in opposite directions (23). This bifurcating replication direction mirrors the reporter orientations that favor DGR editing. To investigate whether DNA replication contributes to this target orientation bias, we measured the repair efficiency of reporters encoded on a plasmid with a p15A origin. Unlike chromosomal replication, replication of a p15A-based plasmid is largely one-way. Although two replication forks form at its origin, one dominant replication fork synthesizes most of the circular plasmid (24) (Fig. 6A). Unlike what’s observed for chromosomal targets (Fig. 5C-D), a single reporter orientation exhibited consistently higher repair efficiency, regardless of reporter location with respect to the origin (Fig. 6B). Crucially, the preferred orientation for DGR targets on this plasmid aligned with the direction of the dominant replication fork. Thus, replication directionality dictates the preferred editing orientation.

**Figure 6.**
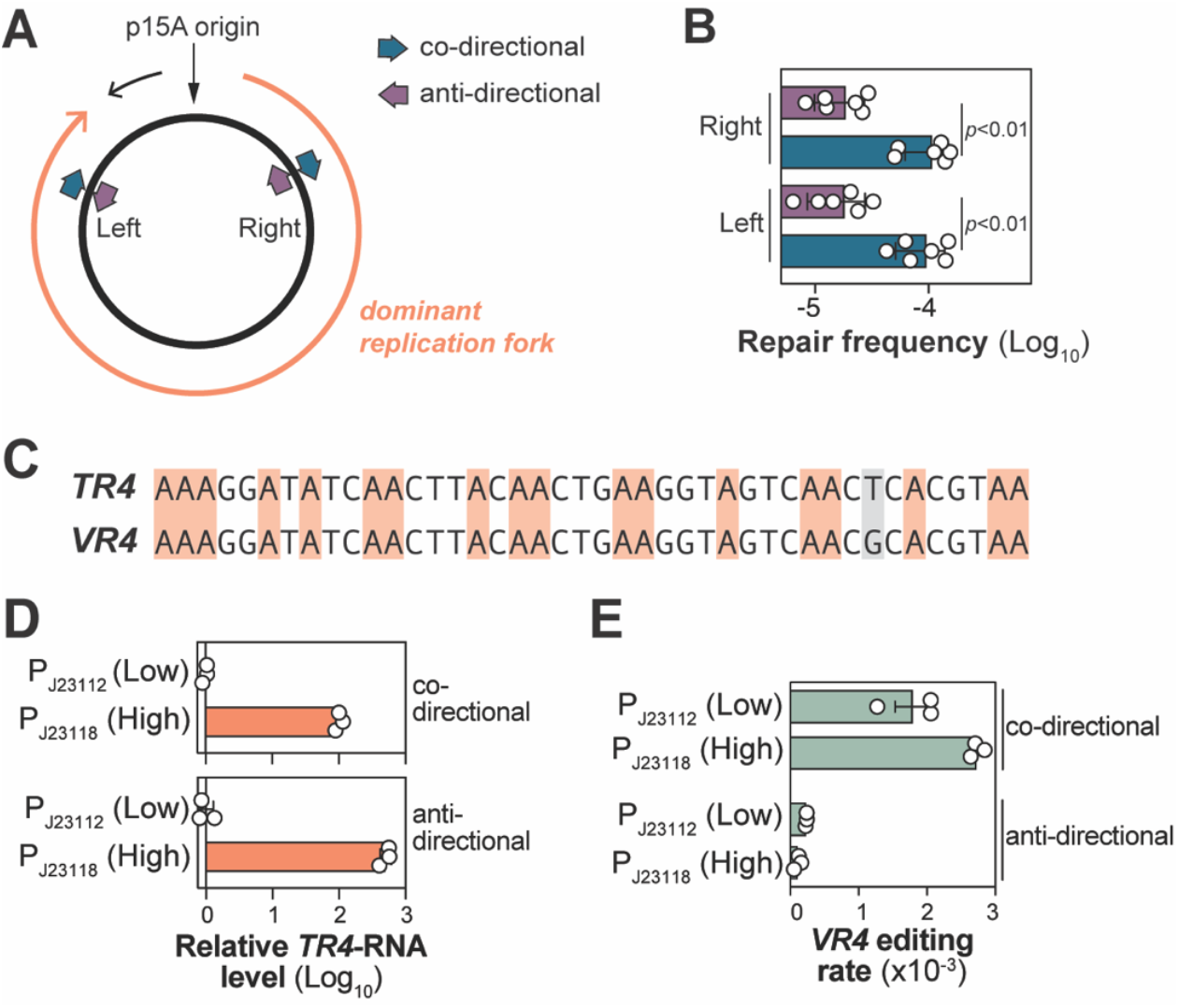
Replication directionality dictates the reporter orientation bias in DGR efficiency. **(A)** Schematic of a plasmid with a p15A origin of replication. The dominant replication fork replicates almost of the entire plasmid (marked by orange arrow). Reporters were inserted in both co-directional and anti-directional orientations relative to the dominant replication fork at the indicated sites, creating pNEB298 (right, anti-directional), pNEB299 (right, co-directional), pNEB300 (left, anti-directional), and pNEB301 (left, co-directional). **(B)** Repair frequency of reporters described in (**A**). Measurements were performed in the *E. coli* MG1655 strain using the kanamycin repair assay, with each p15A-based plasmid co-transformed with pDGR2. Data are the mean of n=6 biological replicates; error bars represent standard deviation. **(C)** Sequences of *TR4* and *VR4* used for deep sequencing analysis of DGR-mediated adenine mutations. Co-transfer of T from *TR4* to replace G in *VR4* (colored grey) was used a signature to distinguish DGR-induced adenine mutations from spontaneous adenine mutations. **(D)** qPCR analysis of reporter expression levels in *E. coli* MG1655. Total RNA was extracted from cells harvested at OD600 ~0.6. The *VR4* reporter, controlled by a weak (P_J23112_) or strong (P_J23118_) constitutive promoter, was integrated at the 291^°^ locus in a derivative of MG1655 strain lacking Δ*sbcB* Δ*mutS*. Reporters were inserted in either co-directional or anti-directional orientations with replication, generating the strains HCL124 (P_J23112_, co-directional), HCL121 (P_J23112_, anti-directional), HCL126 (P_J23118_, co-directional) and HCL123 (P_J23118_, anti-directional). For each orientation, reporter expression levels were normalized to *tufA* and then to the expression level of the P_J23112_. Error bars denote the standard error of the mean for three biological replicates. **(E)** Deep sequencing quantification of DGR-mediated editing rates of the *VRs* described in (**D**). Cells of the indicated *E. coli* strains were grown in LB + 0.2% arabinose until OD600 ~2. Genomic DNA was isolated from harvested cells, and the *VR* regions were PCR amplified for sequencing. The mutagenesis rate of *VR4* was calculated by identifying reads with at least one A-to-N mutation that also the designed G-to-T mutation in the *TR4* region and then dividing it to the total number of reads in that sample. Error bars denote the standard error of the mean for three biological replicates.

### Transcription does not significantly affect DGR editing efficiency

The bacterial chromosomes have evolved to orient highly transcribed genes in the same direction as the movement of DNA replication to prevent head-on collisions between the RNA polymerase and the replisome (25). We therefore considered whether the directional enrichment of stronger promoters could indirectly enhance the transcriptional output of reporters oriented away from *oriC*, which in turns increases their editing efficiency.

To investigate this, we developed a sequencing-based approach to directly measure the rate of A-to-N mutations as a function of the target’s promoter strengths. This method was necessary because altering the expression level of a kanamycin resistance cassette could confound the measurements of resistance. We designed a new reporter (*VR4*) with 18 adenines and drove its expression with two constitutive synthetic promoters: P_J23112_ (weak) and P_J23118_ (strong) (Fig. 6C). To make mutation detection more efficient, we strategically incorporated three key design features: we included many adenines in the high-mutagenesis <u>A</u>AC context (Fig. 3E); we used a strain deleted for *sbcB* and *mutS* (key gene in mismatch repair) (Fig. 4); and we inserted the reporter near *oriC* (291°) (Fig. 5C-D). Furthermore, to control against spontaneous adenine mutation, we also introduced a G-to-T signature mutation into the *TR*-RNA (*TR4*), which was otherwise identical in sequence to *VR4*. The co-transfer of this mutation along with the adenine mutations allowed us to definitively confirm that they were DGR-mediated.

After confirming that these promoters exhibited the expected transcriptional activity (Fig. 6D), we next transformed these strains with a pDGR2 variant to express the corresponding *TR-*RNA (*TR4*) alongside bRT and Avd. We then harvested genomic DNA and quantified the rate of A-to-N mutation by deep sequencing of the reporter locus. Confirming our previous result (Fig. 3E), adenines in the <u>A</u>AC context exhibited significantly higher mutations rates (Fig. S8). Consistent with the results of the kanamycin repair assay, targets oriented co-directional with replication accrued adenine mutations at a higher rate than those in the reverse orientation (weak promoter: 7.6-fold, p-value=0.004; strong promoter: 24.6-fold, p-value < 0.0001) (Fig. 6E). However, for reporters with the same orientation, an orders-of-magnitude difference in transcriptional output only mildly affected the rate of A-to-N mutations (Fig. 6E). Thus, transcription is not a major driver of the observed orientation effects.

## DISCUSSION

Collectively, we demonstrated that it is possible to reconstitute the entire mutagenic retrohoming pathway, spanning from mutagenic cDNA synthesis to target mutagenesis, by transplanting the four core components of the BPP-1 DGR (Fig. 2). This indicates that if other factors are needed to promote mutagenic retrohoming, they are present in *E. coli* and likely other bacteria. Our work also provides a blueprint for using *E. coli* as a heterologous host to study other DGRs from non-tractable organisms or that harbor alternative accessory factors (26).

bRT has a unique error-prone profile that should, in theory, allow it to induce vast sequence diversity into the target gene (2). However, our systematic analysis of cDNA produced by the BPP-1 bRT in *E. coli* revealed a limitation: T-to-C misincorporation dominated most sequence contexts (Fig. 3). This primarily resulted in A-to-G transition in the target, severely restricting the repertoire of target variants that can be generated. We found that the error profile of bRT widened in two specific contexts, showing increased A and G misincorporations in addition to C in the <u>A</u>AC and <u>A</u>C contexts, respectively. In the G<u>A</u>C and C<u>A</u>C contexts, we observed up to 45% G misincorporation (Fig. 3B and E), likely due to the presence of another cytosine at the −2 position (compared to A<u>A</u>C(G) and T<u>A</u>C(A)) (Fig. 3A). This context-dependent switch is particularly striking given that dCTP is about six times more abundant than dGTP in *E. coli* (27).

These ‘deviant’ behaviors—the elevated G misincorporation in the <u>A</u>C context and the increased A misincorporation in the <u>A</u>AC context – are both notably induced by cytosines at the −2 position. This suggests that a G-C DNA-RNA base pair at the −2 position allosterically modifies the template adenine conformation, the bRT active site, or both. Future biophysical and structural studies of this interaction could enable rational engineering of bRT variants with novel mutagenic properties.

Our findings clarify why discrepancies between our results and those from a study using purified BPP-1 bRT *in vitro* (7), which reported a higher overall misincorporation rate (~50% compared to our ~30%) and a preference for adenine over cytosine misincorporation. These differences can be attributed to their use of the native BPP-1 DGR RNA template, where 20 of 23 adenines reside in the AAC context (1)—a context we have found to show higher than average misincorporation rates for both adenine and guanosine (Fig. 3). This interpretation is supported by a recent study that used recombineering to incorporate cDNA generated by BPP-1 bRT into target genes in *E. coli* (28). Through sequencing the target gene (instead of cDNA as we have done here), this work independently identified high mutation rates and altered base preferences in AAC and ACC contexts. Collectively, these findings indicate that the prevalence of AAC motifs in natural TRs is no accident, but rather an evolutionary feature to broaden the range of possible mutations (29).

Our genetic analyses identified ExoI as an endogenous inhibitor of DGR activity in *E. coli* (Fig. 4), and its removal enhanced target editing 15-fold (Fig. 4D-E). Surprisingly, this inhibition does not stem from ExoI-mediated degradation of cDNA-*TR-*RNA (Fig. 4F-H). Furthermore, the observation that increasing cellular ExoI activity beyond a certain point did not further suppress DGR activity suggests an inhibitory mechanism involving interaction with a cellular component (Fig. 4G-H). This finding adds to a growing body of evidence for ExoI having nuclease-independent functions, which remain poorly defined (17,19–22). We propose two models: First, ExoI could sterically hinder cDNA-*TR-*RNA processing or retrohoming by binding the 3’ end of the cDNA, a capacity likely retained by nuclease-deficient variants (30) like Exo^Mut^. Second, ExoI could sequester positive DGR regulators, thereby precluding their involvement.

How cDNA incorporates into the target gene remains the most enigmatic process in DGR-mediated mutagenic retrohoming. Our genome-wide experiments uncovered a prominent *oriC*-centric pattern clearly indicative of DNA replication playing a central role (Fig. 5). Specifically, the higher replication traffic near *oriC*, due to multi-fork replication in *E. coli* (23,31), likely explains the higher target editing efficiency that occurred near the origin. The bidirectional movement of replication forks from *oriC* (32) likely contributes to the preferential editing of target orientation that bifurcates there. Supporting this interpretation, target genes on a unidirectionally replicating plasmid lacked this bifurcation of preferred orientation, showing only a single orientation that aligned with the replication direction (Fig. 6A-B).

Replication likely influences DGR activity by unwinding the target gene to base pair with the cDNA. This mechanism provides a plausible explanation for the observed target orientation bias. Specifically, when the target orientation is co-directional with replication, the cDNA-complementary target sequence is located on the lagging-strand template (Fig. S9A). The discontinuous nature of lagging-strand synthesis keeps this template single-stranded for longer periods, offering an extended window for cDNA binding (33). Thus, DGRs may represent another example of mobile elements that exploit the prolonged single-stranded DNA regions on the lagging strand for insertion into or excision from the chromosome (34).

We attempted to confirm this model, but altering the size of the lagging-strand ssDNA gap had no effect (Fig. S9B). However, we consider these results inconclusive. This is because the cDNA (~70-nucleotide) produced in our experiment is much smaller than the typical ~1-2 kilobase lagging-strand gap (33), so substantial adjustments would be needed to significantly alter efficiency. Based on similar findings in recombineering, such changes may only produce several-fold differences (35), which our current reporter assay may not reliably distinguish.

Additional findings presented here provide independent support for the involvement of DNA unwinding. The three factors identified in our screen that promote DGR editing—RecB, UvrD, and RapA—likely act by unwinding the target (Fig. 4E). RecB and UvrD are helicases that directly unwind DNA (36,37), while RapA acts indirectly by removing transcriptional roadblocks (38). This unwinding mechanism also provides context for the enigmatic stem-loop element downstream of IMH. Our finding suggests that while the stem-loop is not essential for cDNA retrohoming in organisms with high replication activity like *E. coli* (Fig. 2E), it may enhance the efficiency of retrohoming by acting a “molecular wedge” to prolong the single-stranded window of the target. This may also explain why the stem-loop is not universally conserved in DGRs (29). The absence of this stem-loop element in some natural targets may also represent an evolved feature to slow their diversification.

In summary, we used a reconstituted DGR in *E. coli* to identify cellular factors that can alter the efficiency of mutagenic retrohoming by over 15,000-fold (Fig. S10). We envision this system as a powerful tool to better understand how DGRs operate within cells, which will inform efforts to improve its efficiency beyond the current ~1:1000 and widen its utility.

## DATA AVAILABILITY

All sequencing data (raw and processed) are deposited and available from NCBI GEO database under accession numbers: GSE303549, GSE303701, GSE304216, and GSE304215.

## ACKNOWLEDGEMENTS

The authors would like to thank Thomas Evans and Jennifer Ong as early proponents of this work, members of the Lim lab for providing critical feedback, Mehmet Berkmen and Emily McNutt for providing KEIO strains for gene deletions, and David Rudner for feedback on the abstract.

